# Glutamine mimicry suppresses tumor progression through asparagine metabolism in pancreatic ductal adenocarcinoma

**DOI:** 10.1101/2022.10.31.514261

**Authors:** Maria Victoria Recouvreux, Cheska Marie Galapate, Shea Grenier, Edgar Esparza, Yijuan Zhang, Swetha Maganti, Razia Naeem, David A. Scott, Andrew M. Lowy, Hervé Tiriac, Cosimo Commisso

## Abstract

In pancreatic ductal adenocarcinoma (PDAC), glutamine is a critical nutrient that drives a wide array of metabolic and biosynthetic processes that support tumor growth. Despite this established dependency, the targeting of specific enzymes involved in glutamine metabolism is yet to yield any clinical benefit. Here, we have examined the therapeutic potential of 6-diazo-5-oxo-L-norleucine (DON), a glutamine antagonist that broadly inhibits glutamine metabolism. We found that DON treatment significantly blocks PDAC tumor growth and attenuates metastasis. Interestingly, we link the effectiveness of DON in PDAC to asparagine (Asn) metabolism. By inhibiting asparagine synthetase (ASNS), DON significantly reduces intracellular Asn production and Asn supplementation rescues the anti-proliferative effects of DON. We discern that PDAC cells upregulate expression of ASNS as a metabolic adaptation and that modulating ASNS levels can impact DON efficacy. Strikingly, in patient-derived organoids, DON responsiveness is inversely correlated with ASNS expression, a feature that is not observed for other metabolic enzymes targeted by DON. We find that treatment with L-asparaginase (ASNase), an enzyme that catabolizes free Asn, synergizes with DON to impact the viability of PDAC cells. Finally, we identify that a combination therapy of DON and ASNase has a significant impact on metastasis. These results shed light on the mechanisms that drive the effects of glutamine mimicry and point to the utility of co-targeting adaptive responses to control PDAC progression.

## INTRODUCTION

Glutamine is a vital nutrient to tumors as it supports the metabolic and biosynthetic reactions necessary to sustain tumor cell growth. In a process known as glutaminolysis, tumor cells rely on the conversion of glutamine to glutamate, catalyzed by glutaminase (GLS), to replenish TCA cycle intermediates and control redox homeostasis. Glutamine is also critical to the biosynthesis of lipids, nucleotides, and proteins, and can serve as a nitrogen and/or carbon source for the biosynthesis of other amino acids, including asparagine, glutamate, proline, aspartate, alanine, glycine, serine, and cysteine^1,2^. Considering the important role that glutamine has in supporting tumor cell fitness, glutamine metabolism has been recognized as a potential therapeutic target in cancer^3–6^. This could be particularly worthwhile in pancreatic ductal adenocarcinoma (PDAC) since PDAC cells are exquisitely dependent on glutamine for their survival^7,8^. In PDAC, glutamine is the most depleted amino acid relative to adjacent non-neoplastic tissue and contributes the most to TCA cycle metabolites relative to other nutrient sources^9,10^. PDAC cells also require glutaminolysis to drive increases in the NADPH/NADP^+^ ratio that maintain the cellular redox state^8,11^. The extreme reliance of PDAC cells on glutamine may be a unique metabolic feature of these tumors as other tumor types utilize glutamine to a much lesser degree and derive biomass mostly from the catabolism of other amino acids^10,12^.

Despite these established dependencies, the specific targeting of glutaminolysis through GLS inhibition has yet to yield any clinical benefit^13^. In preclinical mouse models of PDAC, the pharmacological inhibition of GLS did not display anti-tumor activity, which was attributed to intratumoral compensatory responses^14^. It is conceivable that the ineffectiveness of GLS inhibitors may reflect that the targeting of GLS alone does not account for the broad metabolic functions that have been ascribed to glutamine in cancer. 6-diazo-5-oxo-L-norleucine (DON) is a glutamine substrate analog that acts as a glutamine antagonist. DON inactivates a variety of glutamine-metabolizing enzymes by competitively and covalently binding to glutamine active sites^15^. As such, DON irreversibly inhibits many metabolic enzymes. DON was originally isolated in the 1950s from *Streptomyces* and some early studies showed anti-tumor activity with moderate toxicities. Recently, low toxicity prodrugs have been developed that are hydrolyzed to DON, the active metabolite, selectively in human tumors^16–19^. The DON prodrugs are themselves inert and while such prodrugs are well-tolerated in humans, they are rapidly metabolized to DON in murine plasma due to carboxylesterase 1 activity^17^; therefore, there is very little benefit to the usage of these prodrugs in the preclinical murine setting. Recent studies have examined how DON might regulate immunometabolism and the immune response in cancer^20,21^; however, little is known about how DON affects the metabolic fitness of the tumor cells themselves and what metabolic adaptations might occur in response to DON as a therapy.

Here, we provide evidence that broadly targeting glutamine metabolism via DON has therapeutic potential in PDAC. Utilizing multiple mouse models, we have found that DON halts PDAC tumor growth and attenuates metastasis. Although DON is best described and utilized as a glutaminolysis inhibitor^22^, we find that DON efficacy in PDAC is specifically linked to asparagine (Asn) availability. Asn is a critical amino acid that supports tumor cell proliferative capacity through its roles in amino acid homeostasis and anabolic metabolism^23^. Moreover, Asn becomes an essential amino acid under conditions where glutamine is limiting^24^. Interestingly, one of the enzymes targeted by DON is asparagine synthetase (ASNS), which uses glutamine as a substrate to synthesize Asn. Although DON was determined to be an ASNS inhibitor in the 1970s^25,26^, its examination as a potential cancer therapy has not considered its effects on ASNS. We find that DON substantially suppresses intracellular Asn production and that Asn supplementation uniquely rescues the anti-proliferative effects of DON. As a metabolic adaptation to DON, both *in vitro* and *in vivo*, we identify that PDAC cells upregulate expression of ASNS. Importantly, suppressing or enhancing the expression levels of ASNS can bolster either sensitization or resistance to DON, respectively. Furthermore, we find that treatment with L-asparaginase (ASNase), an enzyme that catabolizes free Asn into aspartate and ammonium, sensitizes PDAC cells and human PDAC patient-derived organoids (PDOs) to DON. Intriguingly, while combining ASNase with DON treatment *in vivo* does not further accentuate the already profound anti-tumor growth properties of DON alone, the polytherapy has a significant impact on metastatic progression. These results shed light on the metabolic mechanisms driving the antineoplastic effects of glutamine antagonists and point to the utility of co-targeting adaptive responses to such therapies to control PDAC progression.

## RESULTS

### Broadly targeting glutamine metabolism suppresses PDAC progression

To investigate whether broadly targeting glutamine metabolism has the capacity to modulate tumor growth, we initially employed a heterotopic syngeneic mouse model of PDAC that recapitulates the histopathological complexities of the human disease^27^. Murine KPC cells were implanted subcutaneously into the flanks of C57BL/6 mice and either DON or vehicle control was administered to tumor-bearing animals starting when tumors reached a volume of ~130mm^3^. In this setting, DON treatment led to a significant blockade in tumor growth (**Fig. 1a**). To examine the effects of glutamine blockade in the context of endogenous tumors, we further evaluated the effects of DON treatment in an orthotopic syngeneic mouse model of PDAC where KPC cells are surgically implanted directly into the pancreas. Similar to our observations in the heterotopic setting, we found that DON administration led to a substantial and significant decrease in tumor growth relative to the vehicle control (**Fig. 1b**). To interrogate whether the suppressive effects of DON on PDAC tumor growth require an intact immune system, we utilized an orthotopic xenograft mouse model where human PDAC cells were implanted into the pancreata of athymic mice. We found that DON significantly suppressed the growth of orthotopic xenograft tumors derived from either PaTu 8988T cells or primary 779E cells (**Fig. 1c,d**). The concentrations of DON utilized in these various *in vivo* settings had minimal effects on overall animal body weight (**Extended Data Fig. 1a**). Immunohistochemistry of the DON-treated tumors revealed no changes in the extent of apoptosis within the tumors, as measured by cleaved caspase-3 (CC3) staining; however, we detected a decrease in the proliferation markers phospho-Histone H3 (pHis-H3) and Ki-67 (**Fig. 1e and Extended Data Fig. 1b,c**). In addition to the effects on the primary tumors, we also observed a significant decrease in the occurrence of macrometastases to the liver and other organs upon DON treatment using the KPC orthotopic tumor model (**Fig. 1f**). These data indicate that DON has antineoplastic properties in PDAC and that it would be beneficial to examine how DON exerts these anti-tumor effects.

**Fig. 1.**
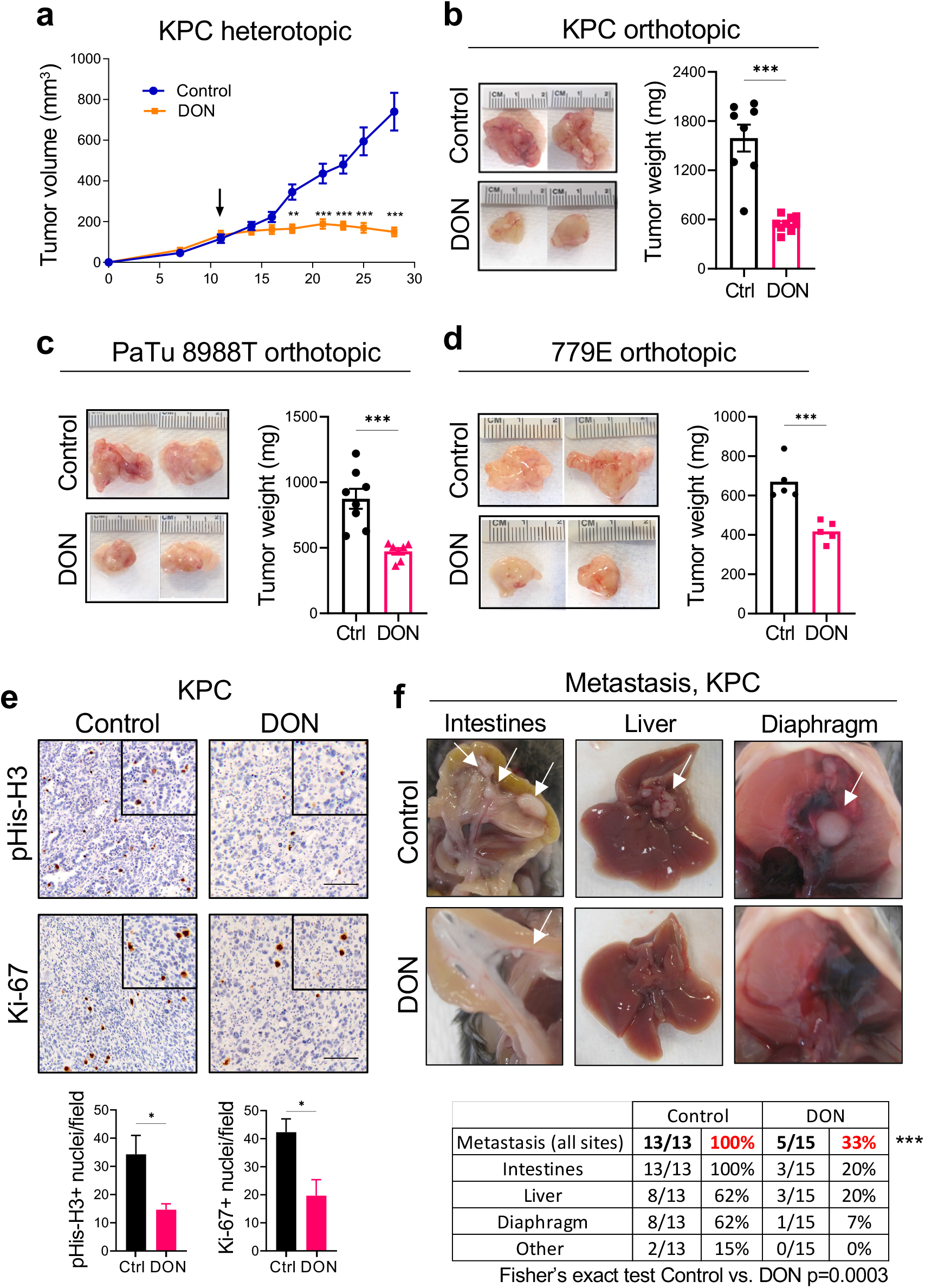
Broadly inhibiting glutamine metabolism suppresses PDAC tumor growth and metastasis. a) Growth of syngeneic KPC tumors implanted subcutaneously and treated with vehicle control (Control) or DON (10mg/kg) for 2 weeks. Tumor volumes were measured with a digital caliper at the indicated timepoints. Data are expressed as mean ± SEM of n=4 mice/8 tumors per group. b) Tumor weights of KPC orthotopic tumors from animals treated with vehicle control (Ctrl) or DON (10mg/kg) for 10 days. Representative tumor images are shown. Data are expressed as mean ± SEM of n=8 or n=9 mice per group (Ctrl, DON). c) Tumor weights of PaTu 8988T orthotopic tumors from animals treated with vehicle control (Ctrl) or DON (10mg/kg) for 3 weeks. Representative tumor images are shown. Data are expressed as mean ± SEM of n=7 mice per group. d) Tumor weights of 779E orthotopic tumors from animals treated with vehicle control (Ctrl) or DON (10mg/kg) for 2 weeks. Representative tumor images are shown. Data are expressed as mean ± SEM of n=8 or n=7 mice per group (Ctrl, DON). e) Immunohistochemical staining of the proliferation markers phospho-Histone H3 (pHis-H3) and Ki-67 in KPC orthotopic tumors treated with vehicle (Ctrl) or DON (10mg/kg). Representative images are shown. Scale bar 100 μm. Quantification of pHis-H3- or Ki-67-positive nuclei/field is shown as mean ± SEM of n=4 tumors per group. f) Characterization of macrometastases in animals with KPC orthotopic tumors treated with vehicle or DON (10mg/kg). Representative images of macrometastases in different tissues are shown. The number of mice with metastases in each organ site was quantified as indicated in the table. n=13 or n=15 mice per group as indicated. Statistical significance was calculated using unpaired two-tailed Student’s t test (a-e) and by Fisher’s Exact test (f). \**P*<0.05, \*\**P*<0.01, \*\*\**P*<0.001.

### DON compromises PDAC cell fitness through Asn metabolism

To better scrutinize the detrimental effects of DON on PDAC cell fitness, we employed an *in vitro* cell-based system. We performed dose-response assays and found that DON treatment severely compromised the proliferative capacity of murine KPC cells, as well as human PDAC cell lines (MIA PaCa-2 and PaTu 8988T) and primary PDAC cells (779E; **Fig. 2a**). DON inhibits a variety of enzymes that utilize glutamine as a substrate, including those involved in metabolic processes such as amino acid biosynthesis, glutaminolysis, hexosamine biosynthesis, and *de novo* pyrimidine and purine biosynthesis^15^. To examine which of these cellular processes might be contributing to the deleterious effects of DON, we took an orthogonal approach and performed metabolite rescue experiments. Surprisingly, although DON has been extensively studied as an inhibitor of glutaminase (GLS or GLS2), cell-permeable α-KG failed to appreciably rescue DON toxicity in these PDAC cell lines (**Fig. 2b and Extended Data Fig. 2a,b)**. We also found that DON antiproliferative effects were not reversed by exogenous addition of N-acetylglucosamine (GlcNAc) or a mixture of nucleosides (Nuc), which would restore hexosamine biosynthesis, and pyrimidine and purine biosynthesis, respectively (**Fig. 2b and Extended Data Fig. 2a,b**). Interestingly, the cell growth defects caused by DON were rescued by supplementation of the growth medium with a cocktail of non-essential amino acids (NEAAs) (**Fig. 2b and Extended Data Fig. 2a,b**). The NEAA cocktail contains several amino acids that are not present in standard DMEM including alanine (Ala), asparagine (Asn), aspartate (Asp), glutamate (Glu), and proline (Pro). We individually assessed each of these amino acids for their ability to rescue proliferative defects caused by DON. Remarkably, only the addition of Asn to the medium had the capacity to rescue PDAC cell growth, at a range of DON concentrations (**Fig. 2c-e and Extended Data Fig. 2c-f**). To determine whether the antiproliferative effects of DON are linked to Asn biosynthesis, we quantified intracellular polar metabolites using gas chromatography/mass spectrometry (GC/MS). We found that in both human and murine PDAC cells, DON treatment significantly reduced intracellular Asn levels (**Fig. 2f**). These observations are in line with previous studies that showed that DON can bind to and inhibit asparagine synthetase (ASNS)^15,25,26^. ASNS catalyzes the synthesis of Asn from Asp via the transfer of the amide group from glutamine to aspartate. Consistent with DON inhibiting ASNS, we found a dose-dependent accumulation of intracellular Asp upon DON treatment (**Extended Data Fig. 2g**). In accordance with its role as a glutaminolysis inhibitor, we also observed significant decreases in Glu and several TCA cycle intermediates, including succinate, fumarate and malate, although the extent to which the TCA cycle metabolites were reduced was variable (**Extended Data Fig. 2h-j**). Altogether, these data demonstrate that DON suppresses Asn biosynthesis and that Asn has the unique ability to rescue the deleterious effects of DON, suggesting that Asn supply and ASNS activity might be distinctively linked to DON effectiveness in PDAC.

**Fig. 2.**
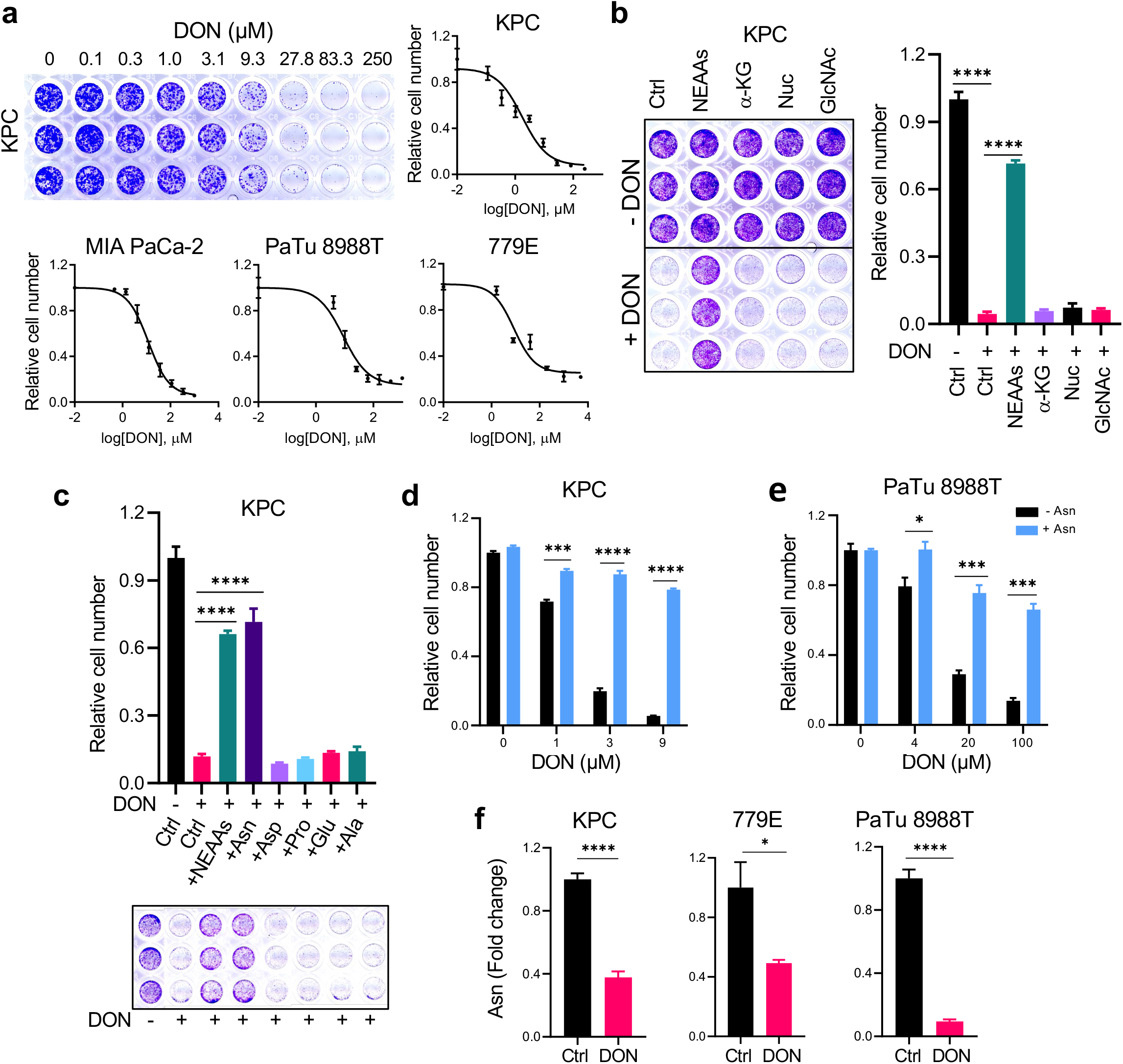
DON treatment suppresses PDAC cell growth through inhibition of asparagine metabolism. a) DON dose response curves for the indicated PDAC cell lines. KPC and MIA PaCa-2 cells were treated for 48h, PaTu 8988T for 24h and 779E for 72h. Relative cell number was quantified by crystal violet staining. Representative images of the crystal violet staining are shown for the KPC cells. Data are presented relative to the untreated control condition and are representative of 3 independent experiments, each performed with 3 replicates per condition. b) Crystal violet assays showing relative cell number of KPC cells treated with or without DON (5μM) and supplemented with the indicated metabolites for 24h. Representative images are shown. Data are presented relative to the untreated control (-DON) for each condition and are representative of 3 independent experiments. Data are expressed as mean ± SEM of triplicate wells. Metabolites assessed were non-essential amino acids (NEAAs), α-ketoglutarate (α-KG), nucleosides (Nuc) and N-acetyl glucosamine (GlcNac). c) Crystal violet assays showing relative cell number of KPC cells treated with or without DON (5 μM) supplemented with either a cocktail of NEAAs or the indicated individual amino acids (0.1mM) for 24h. Representative images are shown. Data are presented relative to the untreated control (-DON) for each condition and are representative of 3 independent experiments. Data are expressed as mean ± SEM of triplicate wells. d), e) Relative number of KPC cells (d) or PaTu 8988T cells (e) treated with DON at the indicated concentrations with or without asparagine (Asn, 0.1mM) supplementation. Data are presented relative to the untreated control condition (-Asn, 0μM DON) and are representative of 3 independent experiments. Data are expressed as mean ± SEM of triplicate wells. f) Quantification of intracellular Asn levels in the indicated PDAC cell lines treated with vehicle control (Ctrl) or DON (10μM for KPC and PaTu 8988T, 1mM for 779E) for 24h. Data are presented relative to untreated control. Data are expressed as mean ± SEM of n=5 wells (KPC, 779E) or n=3 wells (PaTu 8988T). Statistical significance was calculated using One-way ANOVA followed by Dunnet’s multiple comparisons test (b,c) or unpaired two-tailed Student’s t test (d-f). \**P*<0.05, \*\**P*<0.01, \*\*\**P*<0.001, \*\*\*\**P*<0.0001.

### DON effectiveness and ASNS expression are interdependent

Tumor cells exhibit a wide array of adaptive processes that serve to sense and respond to amino acid scarcity. It has been well established that ASNS levels are regulated by nutrient stress^28^; therefore, we surmised that ASNS levels might be affected by DON. Indeed, we found a dose-dependent enhancement of ASNS expression in both human and murine PDAC cells (**Fig. 3a**). This boost in ASNS expression by DON occurred at both the protein and transcript levels and was completely rescued by Asn supplementation of the growth medium (**Fig. 3b,c and Extended Data Fig. 3a,b**). The impact of DON on ASNS expression was also observed *in vivo*, as orthotopic tumors from DON-treated mice displayed significantly higher levels of ASNS protein as measured via western blot and immunohistochemistry (**Fig. 3d-f and Extended Data Fig. 3c**). We next assessed whether ASNS expression levels might modulate the effectiveness of DON. To examine whether reducing ASNS expression has the capacity to sensitize PDAC cells to DON, we knocked down *ASNS* via siRNAs and evaluated DON effects on cell proliferation relative to a non-targeting siRNA. Utilizing this approach, *ASNS* expression levels were partially reduced by approximately 70% (**Fig. 3g and Extended Data Fig. 3d**). Interestingly, ASNS knockdown alone compromised the growth of cells and DON effectiveness was significantly enhanced with partial ASNS depletion (**Fig. 3h,i**). To interrogate whether elevated levels of ASNS might confer resistance to DON treatment, we overexpressed *ASNS* and assessed DON responsiveness. We found that elevated ASNS expression conferred tolerance to DON over a broad range of concentrations (**Fig. 3j,k**). These data indicated that varying ASNS levels can modulate DON efficacy, suggesting that Asn supply could play a role in sensitizing PDAC cells to glutamine antagonism.

**Fig. 3.**
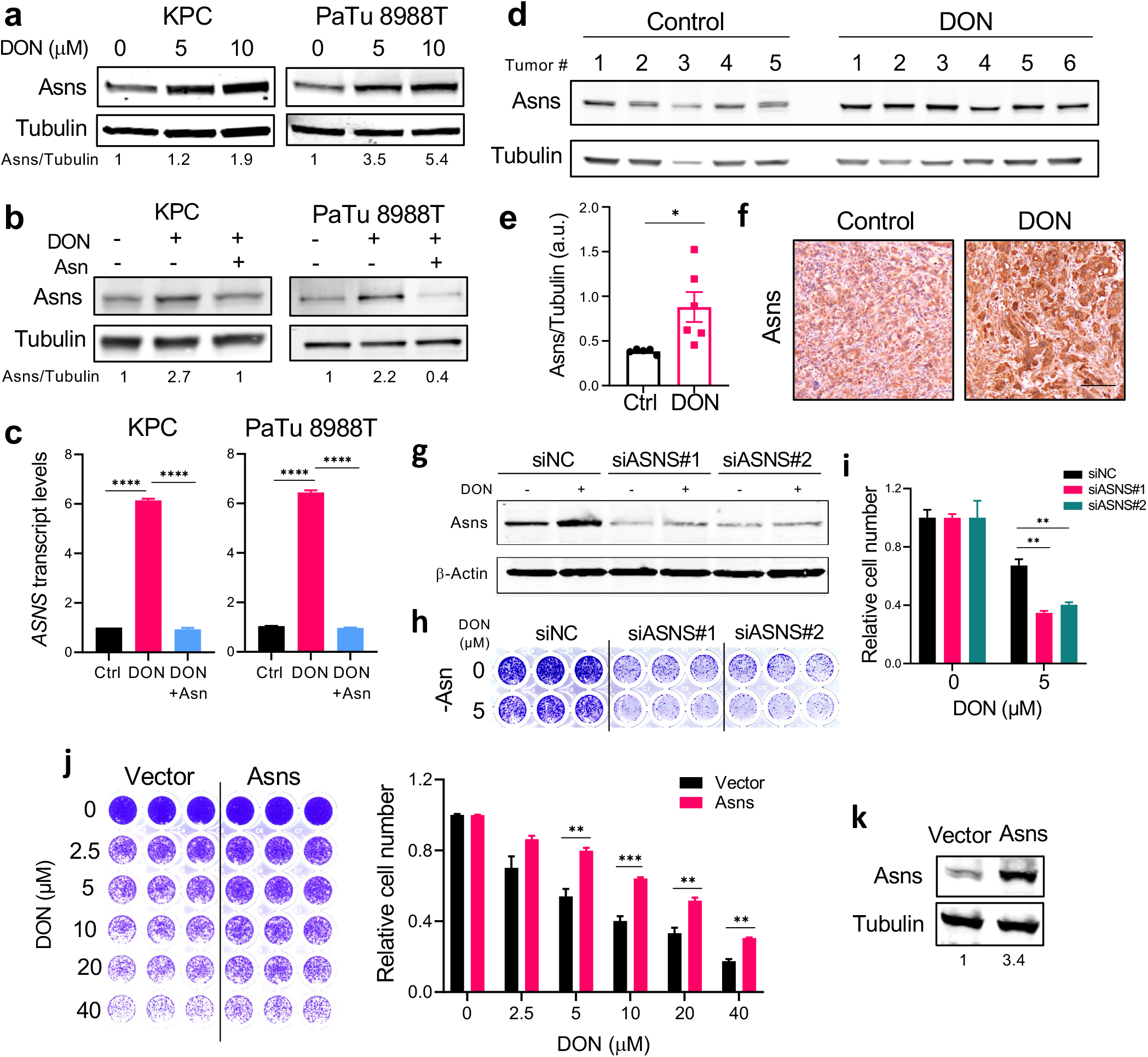
ASNS upregulation is a metabolic adaptation to DON treatment. a) Immunoblot assessing Asns protein levels in KPC and PaTu 8988T cells treated with DON at the indicated concentrations for 24h. Tubulin was used as a loading control. The results are representative of 3 independent experiments. b) Immunoblot assessing Asns protein levels in KPC and PaTu 8988T cells treated with 10μM DON with or without Asn supplementation for 24h. Tubulin was used as a loading control. The results are representative of 3 independent experiments. c) Relative *ASNS* mRNA levels as assessed by qPCR in KPC or PaTu 8988T cells treated with 10μM DON with or without Asn supplementation for 24h. Data are presented relative to untreated control (Ctrl) and are representative of 3 independent experiments. Data are expressed as mean ± SEM of n=3 replicates. d) Immunoblot assessing Asns protein levels in KPC orthotopic tumors treated with vehicle (Control) or DON (10mg/kg) for 2 weeks. Tubulin was used as a loading control. e) Quantification of Asns protein relative to Tubulin in KPC orthotopic tumors from d). Data are expressed as mean ± SEM of n=5 or n=6 tumor samples per group. f) Immunohistochemical staining of Asns protein in KPC orthotopic tumors treated with vehicle control (Control) or DON (10mg/kg). Representative images of n=4 mice per group are shown. Scale bar, 100μm. g) Immunoblot assessing Asns protein levels in KPC cells transfected with non-targeting negative control siRNA (siNC) or two different hairpins targeting *ASNS* (siASNS#1 and siASNS#2) for 24h followed by 5μM DON treatment for 24h. h), i) Relative number of KPC cells after transfection with non-targeting siNC control, or siASNS#1 or siASNS#2 for 24h followed by 5μM DON treatment for 24h. Representative images of n=3 independent experiments (h). Quantification of relative cell number as assessed by crystal violet staining (i) where data is presented relative to untreated control for each hairpin and is representative of 3 independent experiments. Data are expressed ± SEM of n=3 replicate wells. j) Relative number of KPC cells after transfection with empty vector or an ASNS-expressing vector for 24h followed by DON treatment at the indicated concentrations. Quantification of crystal violet staining is shown relative to untreated control for each condition and is representative of 3 independent experiments. Data are expressed ± SEM of n=3 replicate wells. k) Immunoblot assessing Asns protein levels in KPC cells transfected with empty vector or ASNS-expressing vector for 48h. Tubulin was used as loading control. Results are representative of 3 independent experiments. Statistical significance was calculated using One-way ANOVA followed by Tukey’s multiple comparisons test (c) or unpaired two-tailed Student’s t test (e, i and j). \**P*<0.05, \*\**P*<0.01, \*\*\**P*<0.001, \*\*\*\**P*<0.0001.

### ASNase sensitizes PDAC cells to DON and ASNS is a predictive biomarker of DON response

To test the idea that targeting Asn supply might sensitize PDAC cells to DON, we employed L-asparaginase (ASNase), an enzyme that converts Asn to Asp with NH_4_^+^ as a byproduct and is utilized as a therapeutic treatment in acute lymphocytic leukemia (ALL)^29,30^. To mimic the amino acid composition of tumors, we cultured murine and human PDAC cells in physiological levels of Asn and treated with DON in the presence or absence of ASNase. As expected, supplementation with Asn substantially rescued the toxic effects of DON (**Fig 4a,b and Extended Data Fig. 4a**). Strikingly, DON co-treatment with ASNase led to synergistic effects on cell fitness (**Fig 4a,b and Extended Data Fig. 4a**). ASNase on its own had no effect on cell growth, as it does not prevent intracellular Asn biosynthesis by ASNS. Synergy was computed using the coefficient of drug interaction (CI). To further assess the relationship between DON and ASNase, we employed previously characterized human PDAC patient-derived organoids (PDOs)^31,32^. Single agent and combination treatments of DON and ASNase in a panel of six PDOs revealed a broad range of sensitivities to DON as a single agent (**Fig. 4c**). The PDOs were cultured in the presence of physiological levels of Asn; hence it was not surprising that ASNase alone had only marginal effects or no impact on PDO viability. Significant synergy for the DON + ASNase combination was observed in 5/6 PDOs, as indicated by a CI computation of <0.7 (**Fig. 4c**). The PDOs that we used had already been characterized at the transcriptomic level via RNASeq^32^; therefore, we discerned molecular features that related to drug responsiveness. Heatmap analysis of normalized gene expression for the known DON targets strikingly demonstrated that ASNS is the only DON-targeting enzyme for which heightened expression correlates with dampened DON response (**Fig. 4d**). This relationship between DON and ASNS was confirmed utilizing a Spearman’s rank correlation coefficient analysis for all the known DON targets, and surprisingly, this assessment indicated that the expression of well-established DON targets such as GLS, GLS2 and GFAT (encoded by GFPT1) did not correlate with DON sensitivity (**Fig. 4e and Extended Data Fig 4b-i**). Altogether, these data demonstrate that in PDAC cells ASNS expression levels are inversely correlated with DON efficacy and that ASNase can synergize with DON to reduce cell fitness.

**Fig. 4.**
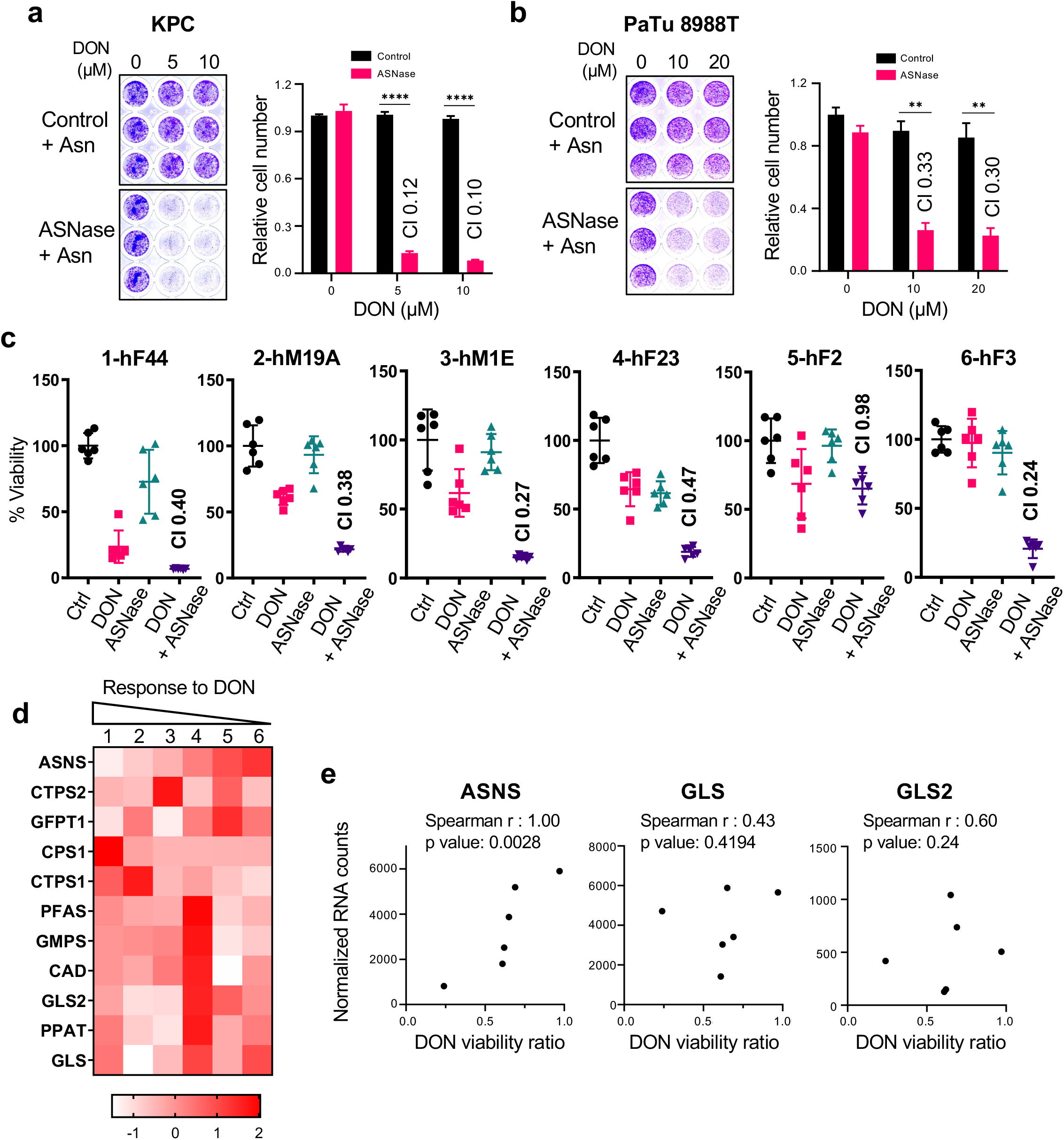
ASNase synergizes with DON to suppress PDAC cell growth. a), b) Relative number of KPC cells (a) or PaTu 8988T cells (b) treated with the indicated doses of DON in combination with L-asparaginase (ASNase, 0.5U/ml) for 24h. Cells were stained with crystal violet, representative images are shown. Quantification of crystal violet staining is shown relative to the untreated control and is representative of 3 independent experiments. The coefficient of drug interaction (CI) was calculated for each DON concentration (shown in graph). Data are expressed ± SEM of n=3 replicate wells. c) Cell viability was assessed in 6 PDAC patient-derived organoids (PDOs) treated with vehicle (Ctrl), 50μM DON, 0.3 U/ml ASNase, or combination of DON + ASNase. The CI is indicated in the graphs. Data are presented relative to untreated control and are representative of 2 independent experiments, with 5 replicates each. Data are expressed as mean ± SEM of n=5 replicate wells. d) Heatmap analysis of normalized gene expression (RNA-seq) of known DON-targeting enzymes in PDAC patient-derived organoids (PDO) used in c. e) Correlation analysis between normalized gene expression of the indicated DON-targeting enzymes and DON viability ratio, used as measurement of PDO responsiveness to DON treatment. Statistical significance was calculated using unpaired two-tailed Student’s t test (a,b) or Spearman correlation coefficient (e). \*\**P*<0.01, \*\*\*\**P*<0.0001.

### DON and ASNase combination therapy restrains PDAC progression

To investigate the therapeutic potential of combining DON with ASNase in PDAC, we employed an orthotopic xenograft mouse model where endogenous pancreatic tumors were derived from human PaTu 8988T cells. PDAC cells were implanted directly into the pancreata of athymic mice and approximately 20-23 days post-implantation when tumors were palpable, animals were treated with either vehicle control, ASNase alone, DON alone or DON + ASNase for two weeks, at which point tumors were extracted for analysis. Weights of the primary tumors revealed that ASNase alone did not affect tumor size, while DON alone, as expected, resulted in a significant diminishment in tumor growth (**Fig. 5a**). The combination of DON + ASNase did not lead to a further reduction in weight of the primary tumors (**Fig. 5a**). Consistent with this finding, markers for proliferation and cell death in DON-treated tumors were not modulated by the combination treatment (**Extended Data Fig. 5a-b**). In ALL therapy, ASNase is utilized to deplete circulating Asn; therefore, to confirm ASNase activity in our orthotopic xenograft mouse model, we quantified Asn plasma levels. Indeed, we found that ASNase treatment led to an extensive depletion of circulating Asn (**Fig. 5b**). In contrast, however, intratumoral depletion of Asn in the orthotopic pancreatic tumors was less impacted (**Fig. 5c**). These data indicate that while ASNase efficiently depletes circulating Asn, tumor Asn levels are less affected. We reasoned that with the diminishment of circulating Asn, the anti-tumor effects of DON might be accentuated at the level of metastatic progression. Indeed, with the combination of DON and ASNase we observed a substantial decrease in the number of animals displaying macrometastases to the intestines, liver, and diaphragm (**Fig. 5d,e and Extended Data Fig. 5c**). Altogether, our observations suggest that while ASNase treatment may not further enhance the robust anti-tumor effects of DON alone in the primary PDAC tumors, a DON + ASNase polytherapy may provide a combinatorial benefit in controlling metastasis.

**Fig. 5.**
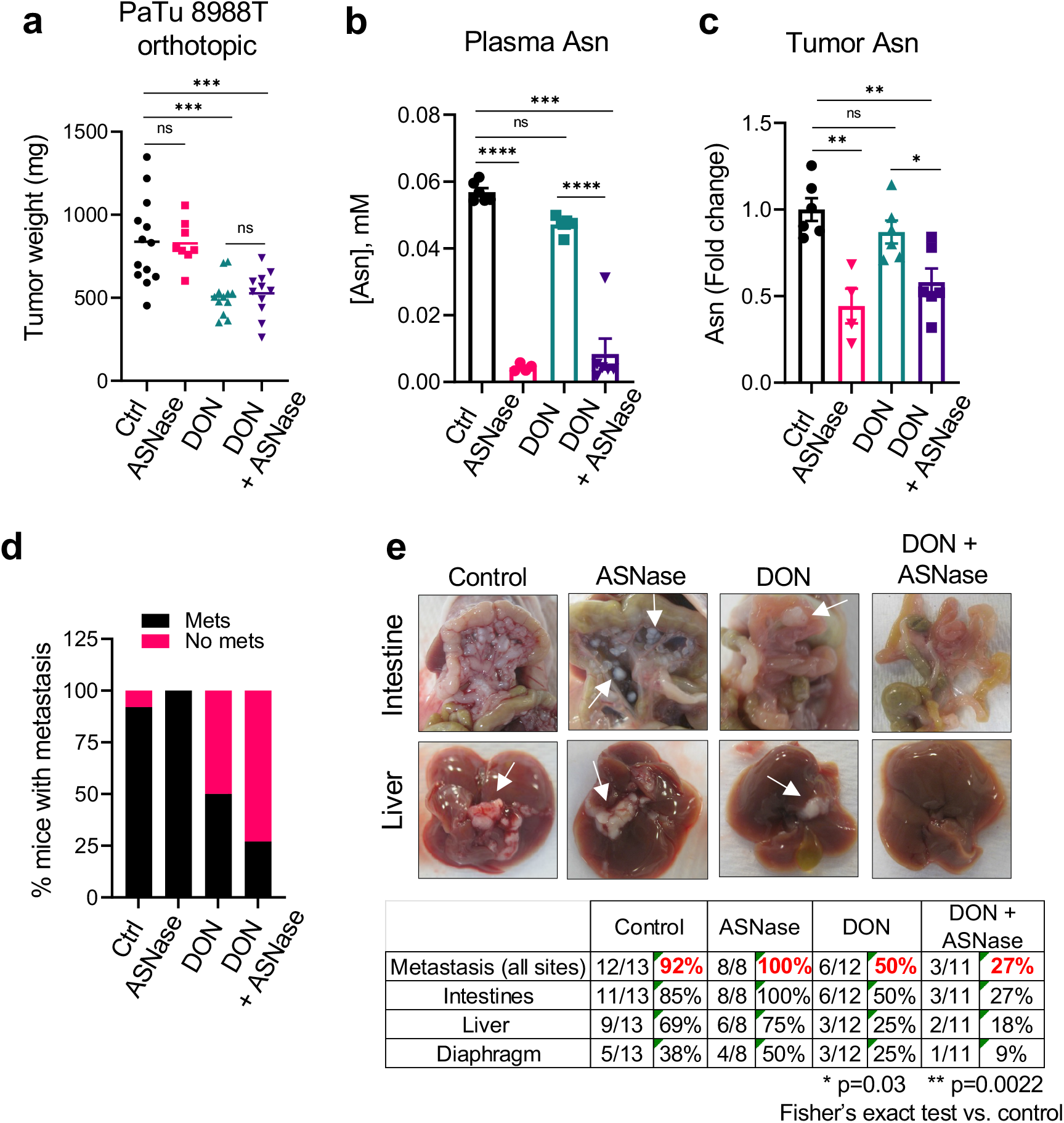
DON and ASNase combination therapy suppresses PDAC metastasis. a) Tumor weights of PaTu 8988T orthotopic tumors from animals treated with vehicle (Ctrl), ASNase (60U/mouse), DON (5-10mg/kg), or combination of DON + ASNase for 3 weeks. Data are expressed as mean ± SEM of n=13, n=8, n=12, or n=11 mice combined from 2 different cohorts. b) Asn levels quantified by GC-MS in plasma of mice bearing PaTu 8988T orthotopic tumors treated as indicated. Data are expressed as mean ± SEM of n=6, n=4, n=5, or n=6 mice per group. c) Quantification of intratumoral Asn levels in PaTu 8988T orthotopic tumors treated as indicated. Data are presented relative to no treatment control and quantification was calculated as amount of Asn per mg of tumor tissue. Data are expressed as mean ± SEM of n=6, n=4, n=6, or n=6 mice per group. d) Quantification of the percentage of mice with PaTu 8988T-derived primary orthotopic tumors that presented with macrometastases. n=13, n=8, n=12, or n=11 mice per group. e) Characterization of macrometastases in animals with PaTu 8988T orthotopic tumors treated as indicated. Representative images of macrometastases in the different tissues are shown. The white arrows point to macrometastases. The number of mice with macrometastases in each organ site was quantified as indicated in the table. n=13, n=8, n=12, or n=11 mice per group. Statistical significance was calculated using one-way ANOVA followed by Tukey’s multiple comparisons (a-c) or Fisher’s exact test (e). \**P*<0.05, \*\**P*<0.01, \*\*\**P*<0.001, \*\*\*\**P*<0.0001.

## DISCUSSION

Our studies have elucidated the tumor cell intrinsic effects of broadly inhibiting glutamine metabolism in PDAC. We delineate how glutamine mimicry by DON profoundly suppresses cellular fitness through the regulation of Asn production, and how DON treatment leads to significant abrogation of tumor growth and metastasis in mouse models of PDAC. Previous studies have mainly focused on tumor cell extrinsic effects of DON by investigating how DON and DON-based prodrugs impact the tumor ecosystem through modulating stromal cell function. In an immunoresponsive syngeneic mouse model in which heterotopic tumors were derived from murine MC38 colon cancer cells, JHU083, a DON-based prodrug, enhanced endogenous anti-tumor immune responses by boosting tumor infiltration and activation of CD8^+^ T cells^20^. In this MC38 model, JHU083 treatment in *Rag2^−/−^* mice, which do not produce mature B or T lymphocytes, failed to affect tumor growth rates and animals succumbed to disease in a similar time frame as untreated wild-type mice. In contrast, our experiments utilizing athymic nude *Foxn1^nu^* mice, which are unable to produce T cells, revealed that DON significantly suppresses the growth of orthotopic xenograft tumors derived from human PDAC cells. These data indicate that tumor growth control by DON treatment in our immunocompetent PDAC mouse models is not at all mediated by T cells, or that the contribution of the endogenous immune responses in these settings is minimal. DON has also been implicated in regulating extracellular matrix (ECM) remodeling in an orthotopic cancer-associated fibroblast (CAF) co-implantation model of PDAC^21^. In this model, intrapancreatic co-implantation of a high ratio of CAF:KPC cells (9:1) allows for the optimal evaluation of CAF-secreted ECM molecules, such as collagen and hyaluronan (HA). It was found that in these tumors, DON decreases HA production likely through its inhibition of GFAT1, a known DON target that regulates UDP-GlcNAc synthesis. Interestingly, in this model, the effects of DON on tumor growth were also attributed to CD8^+^ T cells, as the efficacy of DON disappeared in CD8 knockout mice. In our orthogonal rescue assays, providing exogenous GlcNAc did not rescue the anti-neoplastic effects of DON, as well, in our human PDO evaluations, GFAT1 expression levels (encoded by the *GFPT1* gene) did not correlate with DON responsiveness. Further scrutinization into the multipronged effects of DON will be required to evaluate the specific contributions of each of the DON targets to *in vivo* tumor growth and the extent to which tumor-stromal interactions control DON responsiveness in preclinical models and patients.

The current findings indicate that a combination approach utilizing DON together with ASNase might represent a novel therapeutic modality to control metastasis in PDAC. ASNase treatment is a key chemotherapy that is employed in ALL, the most common pediatric cancer. ASNase effectiveness in ALL has been attributed to the fact that lymphoblasts lack expression of ASNS, thus they depend on the uptake of circulating Asn for their survival^33^. The potential of ASNase as a therapy in solid tumors has been limited; however, in PDAC preclinical models, combination therapies including ASNase have been demonstrated to have efficacy at controlling tumor growth. In a syngeneic orthotopic mouse model of PDAC, ASNase in combination with phenformin, a metformin-related electron transport chain (ETC) inhibitor, significantly reduced tumor growth^34^. This important finding is directly linked to ETC inhibition effectively impairing *de novo* Asn biosynthesis, and the ability of exogenous Asn to restore proliferation upon ETC inhibition^34^. Interestingly, in human PDAC cells, the *in vitro* depletion of Asn by ASNase has been shown to enhance Erk1/2 phosphorylation, and the deleterious proliferative effects of *ASNS* knockdown were further enhanced by MEK inhibition^35^. Importantly, the *in vivo* combination of ASNase and MEK inhibition suppressed the growth of orthotopic PDAC tumors to a greater extent than either of the single agents alone^35^. Apart from our examination of combining DON and ASNase *in vivo*, little is known about combination therapies with DON that may improve its anti-tumor effects in PDAC. However, in preclinical mouse models of glioblastoma (GBM), DON in combination with a calorie-restricted ketogenic diet (KD) resulted in reduced tumor growth and enhanced survival^36^. The mechanistic underpinnings of how the KD improves the effectiveness of DON in GBM remain unclear and whether similar diet modifications synergize with DON in PDAC is an interesting area for future interrogation.

The determination that ASNS expression levels negatively correlate with DON sensitivity in human PDOs suggests that ASNS might be a predictive biomarker of DON effectiveness in PDAC. We demonstrated that indeed modulating ASNS expression levels in PDAC cells has the capacity to control DON sensitivity or tolerance. Interestingly, this observation was unique to ASNS among the known DON target enzymes, as expression of well-studied targets such as GLS and GLS2 did not correlate in this way. Very little is known about what might dictate patient stratification for DON-based prodrugs that are currently in clinical trials^37^. Interestingly, in atypical teratoid/rhabdoid tumors (ATRT), which is a rare and aggressive cancer of the brain and spinal cord in infants, metabolic profiling revealed a unique dependence on glutamine for survival in ATRT cell lines with high *MYC* expression^38^. DON selectively suppressed proliferative capacity and induced cell death in these Myc^high^ ATRT cells and extended survival in an ATRT Myc^high^ orthotopic mouse model^38^. These findings are in accordance with a DON-based prodrug that suppressed proliferation and induced cell death in *MYC*-driven medulloblastoma cell lines, and increased survival in an orthotopic xenograft mouse model^39^. These links between Myc and sensitivity to DON are a direct result of the established role that Myc has in regulating metabolic reprogramming. Myc directly reprograms cellular transcriptional outputs that drive mitochondrial glutaminolysis. This transcriptional rewiring by Myc leads to the enhanced cell viability and TCA cycle anaplerosis, while concomitantly creating a dependency on glutamine catabolism^40^. Depriving Myc-driven cancers, such as neuroblastomas, selectively leads to apoptosis through ATF4-dependent mechanisms^41^. It is not yet clear whether ASNS plays a role in modulating the sensitivity of Myc-driven neuroblastomas to glutamine deprivation. Moreover, it remains to be explored whether ASNS expression levels represent a predictive biomarker of response to glutamine metabolism-based therapies in malignancies other than PDAC.

## MATERIALS AND METHODS

### Cells and cell culture conditions

MIA PaCa-2 cells were obtained from the American Type Culture Collection (ATCC); PaTu 8988T cells were purchased from Elabscience. 779E are epithelial cells established from a moderate-to-poorly differentiated patient derived tumor by A.M. Lowy^42^. KPC cells originated from *Pdx1-Cre; LSL-KRas^G12D/+^; LSL-Tp53^R172H/+^* mice were kindly provided by Dr. Robert H. Vonderheide^27^ and the parental KPC cell line (KPC4662) was subcloned to generate KPC#65 and KPC#74 clones that were used in this study. All cell lines were cultured in DMEM supplemented with 10% FBS and 100 units/mL penicillin/streptomycin under 5% CO_2_ at 37C and were routinely tested for mycoplasma contamination by ABM’s PCR Mycoplasma detection kit. The following inhibitors were used in this study: 6-diazo-5-oxo-L-norleucine (DON) (Sigma) and L-Asparaginase (Prospec bio). For rescue experiments cells were treated with MEM-NEAA (Thermo Fisher, 0.1mM); dimethyl-2-oxo-glutarate (α-KG, Sigma, 5.5mM); EmbryoMax Nucleosides 100X (Millipore, 2X) and N-acetyl glucosamine (GlcNAc, Sigma, 10mM). Individual non-essential amino acids, Asn, Asp, Glu, Pro, Ala were obtained from Sigma and used at 0.1mM.

### Animal studies

C57BL/6J female mice and NU/J athymic nude female mice aged between 5 and 6 weeks old were purchased from The Jackson Laboratories and Charles River, respectively. All animals were housed in sterile caging and maintained under pathogen-free conditions. All experimental procedures in mice were approved by the Institutional Animal Care and Committee (IACUC) of Sanford Burnham Prebys Medical Discovery Institute (approval number 20-073).

#### Subcutaneous tumor model

500,000 KPC cells suspended in 100 μL of 1:1 PBS: Matrigel were subcutaneously injected in both lower back flanks of syngeneic C57BL/6J mice under isoflurane anesthesia. Tumor growth was monitored by volume measurement with a digital caliper. When tumors reached a volume of 150-200 mm^3^, mice were randomly assigned into groups and received the following treatments via i.p. administration: control group (vehicle, sterile water); DON-treated group (1, 5, or 10mg/kg as indicated). Mice were injected every other day for 2 weeks. Tumor tissue was collected in 10% formalin for immunostaining and a portion of the tissue was snap-frozen in liquid nitrogen for qPCR analysis.

#### Orthotopic tumor model

12,500 murine or human PDAC cells were injected (50 μL 1:1 PBS: Matrigel) into the tail of the pancreas via a small abdominal incision in the left flank of anesthetized mice. Murine KPC cells were injected into C57BL/6J female mice; and human 779E and PaTu 8988T cells were injected into Nu/J athymic nude mice. Tumor growth was monitored by palpation under isoflurane anesthesia. When tumors were palpable (12 to 20 days after injection depending on the cell line) mice were randomized and treated as indicated: control group, (sterile water, i.p.) and DON-treated group (10mg/kg DON, i.p.). Mice were injected every other day for 2 weeks. For DON + L-Asparaginase (ASNase) combination treatment, PaTu 8988T tumor-bearing mice were randomized into 4 groups: Control (sterile water, i.p.), DON (5-10 mg/kg, i.p.), L-Asparaginase (3Units/g, i.p.) and DON + ASNase (5-10mg/kg DON + 3U/g ASNase, i.p.). Mice were injected 3 times per week for 2 weeks. Tumors were weighed and collected in 10% formalin for histology and IHC or snap-frozen in liquid nitrogen for metabolite quantification. All mice were thoroughly examined for spontaneous metastasis in the peritoneal cavity (small and large intestines, liver, kidney), diaphragm and lungs. The number of mice presenting metastasis in each of the sites was compared among treatment groups.

### PDAC patient-derived organoid growth

Establishment and DNA and RNA sequencing of human PDAC patient-derived organoids (PDO) used in this study were reported previously^32^. Human organoids were cultured as described previously^32^. Briefly, cells were resuspended in 100% growth factor–Reduced Matrigel (Corning) and plated as domes overlaid with Human Complete Feeding Medium (hCPLT): advanced DMEM/F12, 10 mmol/L HEPES, 1× Glutamax, 500 nmol/L A83– 01, 50 ng/mL hEGF, 100 ng/mL mNoggin, 100 ng/mL hFGF10, 0.01 mmol/L hGastrin I, 1.25 mmol/L N-acetylcysteine, 10 mmol/L Nicotinamide, 1X B27 supplement, 10% (vol/vol) R-spondin1–conditioned medium, 50% (vol/vol) Afamin/Wnt3A conditioned media^43,44^. PDO viability was evaluated with CellTiter Glo luminescent assay (Promega), following manufacturer instructions.

### Pharmacotyping of organoids

Organoids were dissociated into small fragments and approximately one thousand viable cells were plated per well of a 384 well plate in 20 μL 10% Matrigel/ human complete organoid media. Therapeutic compounds were added 24 hours after plating, after the reformation of organoids was visually verified. Each condition was tested in six replicates. Compounds were dissolved in DMSO or PBS. After 5 days, cell viability was assessed using CellTiter-Glo as per the manufacturer’s instruction (Promega) on a plate reader (BioTeck Synergy H1).

### Immunohistochemistry

PDAC tumors were fixed in 10% formalin. Fixed tissue was embedded in paraffin and sectioned by the SBP Histology Core. Immunohistochemistry was performed as previously described^45^. Briefly, antigen retrieval was performed by microwave-heating in 10mM sodium citrate (pH 6) and endogenous peroxidases were quenched in 3% hydrogen peroxide. Sections were blocked in 2% BSA, 10% goat serum in PBS for 1 hour at room temperature and incubated with primary antibodies diluted in 2% BSA/PBS overnight at 4°C. After washes, sections were incubated with biotinylated goat anti-rabbit secondary antibody for 1.5 hours at room temperature followed by incubation with the VECTASTAIN Elite ABC HRP Kit (Vector Labs) and the DAB HRP Substrate Kit (Vector Labs). Nuclear counterstaining was performed by hematoxylin staining. Images were captured with a brightfield Olympus CX-31 microscope coupled with INFINITY camera and INFINITY capture software (Lumenera). The following primary antibodies and dilutions were used: ASNS (1:500, ProteinTech, 14681-1-AP), Cleaved caspase 3 (1:1000, CST, 9664), Ki-67 (1:400, ThermoFisher, MA5-14520) and phospho Histone H3 (1:200, CST, 9701).

### Polar metabolites extraction and quantification

For quantification of polar metabolites in cell lysates, KPC74, PaTu 8988T and 779E cells were seeded in 6 well plates and treated with DON for 24h at the indicated concentrations. Experiments were run in 3-5 replicate wells and extra wells were used for cell counting and protein quantification for normalization. After treatment cells were washed 3X with cold PBS and extracted with ice-cold 50% Methanol/ 20uM L-Norvaline followed by addition of chloroform, mixing by vortex and centrifugation at 14000 RPM for 5min at 4°C. The top layer was dried using a Speedvac, derivatized and analyzed using gas chromatography–mass spectrometry (GC–MS) to quantify small polar metabolites as previously described^46^. For quantification of tumor Asn, PaTu 8988T tumors treated with vehicle, DON, ASNase or combination of DON + ASNase were collected and flash-frozen in liquid nitrogen. 20– 30 mg of tissue samples was transferred to 2-mL tubes containing 2.8 mm ceramic beads (Omni International) and 0.45 ml ice-cold 50% methanol/ 20 mM L-norvaline. Tubes were shaken (setting 5.5) for 30 s on a Bead Ruptor 12 (Omni International), and quickly placed on ice. Samples were centrifuged at 15,000 x g for 10 minutes at 4°C. The supernatant was then transferred to a new tube, mixed with chloroform, and processed in the same way as described for the cell lysates.

### RNA extraction and quantitative real time PCR (qPCR)

Total RNA was extracted from cells or tumor tissue with the PureLink RNA Mini Kit (Thermo Fisher Scientific). For quantitative real time PCR (qPCR), cDNA was synthesized from 1000 ng of total RNA using the High-Capacity cDNA Reverse Transcription Kit (Thermo Fisher Scientific). cDNA samples were diluted 1:10 in RNAse free water. qPCR was performed in triplicates with the SYBR Premix Taq II master mix (Takara) and specific primers on the LightCycler 96 (Roche). Relative target gene expression was determined by comparing average threshold cycles (CT) with that of housekeeping genes (*18s* or *Rpl13*) by the delta-delta CT method. The following primers were used in this study: mASNS-F: CCTCTGCTCCACCTTCTCT; mASNS-R: GATCTTCATCGCACTCAGACA; hASNS-F: GAGTCAGACCTTTGTTTAAAGCA; hASNS-R: GGAGTGCTTCAATGTAACAAGAC; 18s-F: GTAACCCGTTGAACCCCATT; 18s-R: CCATCCAATCGGTAGTAGCG; hRPL13-F: GTTCGGTACCACACGAAGGT; hRPL13-R: TGGGGAAGAGGATGAGTTTG

### Immunoblotting

Cells were lysed in RIPA buffer (10mM Tris-HCl [pH 8.0], 150mM NaCl, 1% sodium deoxycholate, 0.1% SDS, 1% Triton X-100) with protease and phosphatase inhibitors (Roche). Protein concentrations were measured using the DC Protein Assay Kit (Bio-Rad). 15-20 μg protein samples were run in SDS–PAGE followed by protein transfer using Mini Gel Tank and Mini Blot Module (Life Technologies). Immunoblotting was detected using near-infrared fluorescence (LI-COR) and the Odyssey CLx imager (LI-COR). The following primary antibodies were used: ASNS (Proteintech, 1:500), Tubulin (Sigma T6074, 1:10000), and β-actin (Sigma, A1978, 1:20000). The band fluorescence intensities were quantified with ImageStudio Lite software (LI-COR).

### siRNA and plasmid transfection

Cells were transfected in 6-well plates with Lipofectamine RNAiMax Transfection reagent (ThermoFisher Scientific) and murine Asns-targeting siRNA (s7704 and s7705, ThermoFisher Scientific) or negative control siRNA (siNC) at a final concentration of 25nM following manufacturer’s protocol. After 24h, transfected cells were trypsinized and seeded into 96-well plate format for DON treatment experiments. ASNS overexpression was achieved via transfection of a murine ASNS expression vector under a CMV6 promoter (MC200523, OriGene). Cells were transfected in 6-well plates with Lipofectamine 2000 (ThermoFisher Scientific) and 2.5ug of DNA (ASNS or empty vector). After 3-7 days selection with 2mg/ml G418, cells were seeded in 96 well for DON treatment.

### Cell number quantification by crystal violet staining

Cells were seeded in complete culture media at a density of 5000 to 10,000 cells per well in 96-well format. 24 to 48h after seeding, cells were treated with variable concentrations of DON as indicated. For co-treatments of DON and ASNase or rescue metabolites, compounds were added at the same time for 24h. At the indicated timepoints, cells were fixed with 4% paraformaldehyde and stained with 0.5% crystal violet, thoroughly washed, and dried overnight. The crystal violet plates were scanned, and relative cell number was calculated using quantification of the stained area with ImageJ software. Three replicates per condition were used in each experiment. The coefficient of drug interaction (CI) was calculated as: CI = AB/(A×B). Where AB is the ratio of the combination groups to control group; A or B is the ratio of the single agent group to control group.

### Statistical analysis

All graphs and statistical analysis were done using GraphPad Prism software version 9 (GraphPad). Results are shown as means ± standard error of the mean (SEM). Statistical significance was determined by the unpaired two-tailed Student’s t test with Welch’s correction, when appropriate. Comparison of more than 2 groups was done by one-way ANOVA followed by Tukey or Dunnet test for multiple comparisons. *P* values lower than .05 were considered statistically significant. The number of mice with metastases was compared with the Fisher’s Exact test. (\**P*<0.05, \*\**P*<0.01, \*\*\**P*<0.001, \*\*\*\**P*<0.0001).

---

**Extended Data Fig. 1.**
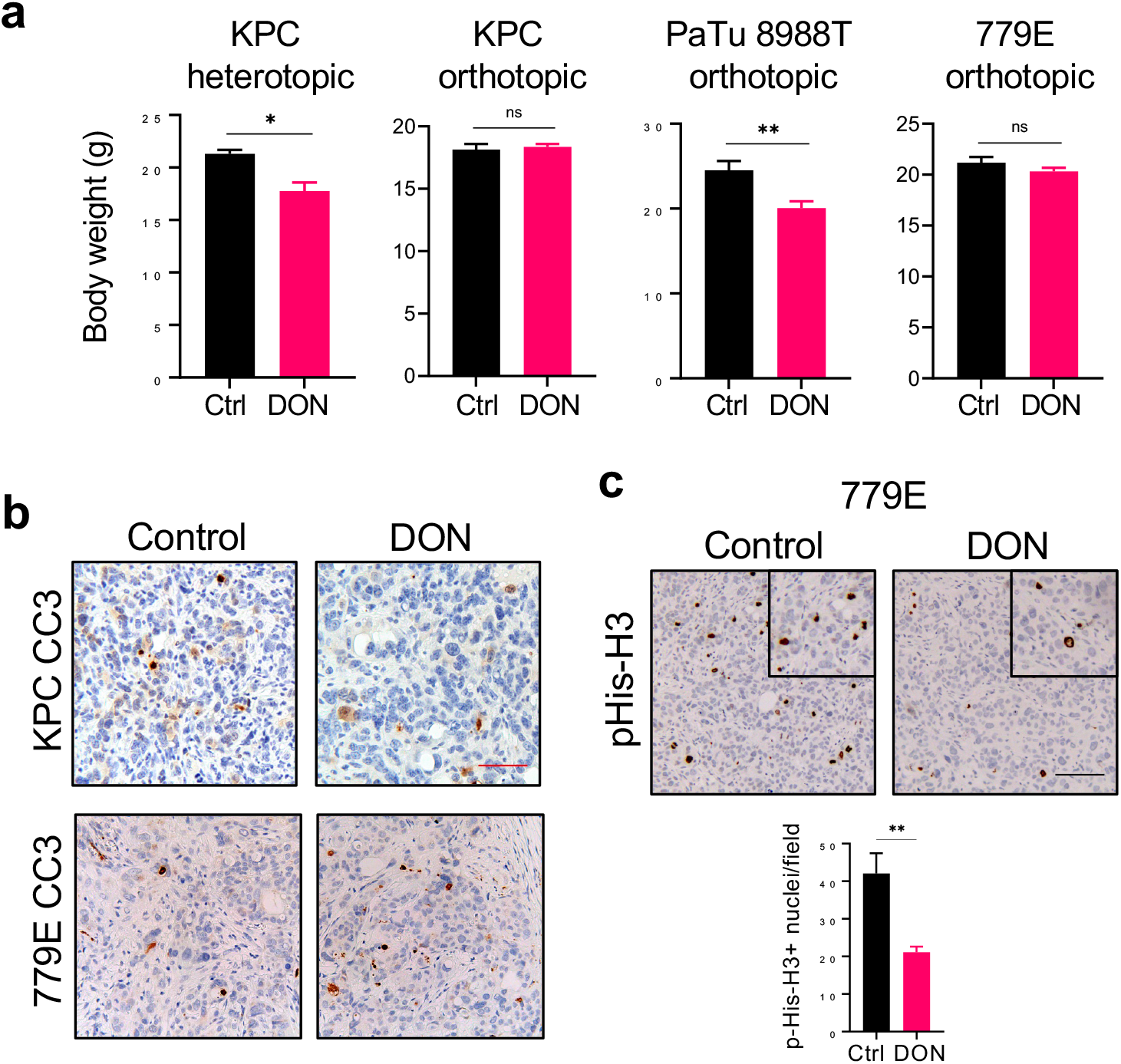
DON treatment minimally affects animal body weight and tumor cell death, but significantly impacts tumor cell proliferation. a) Body weights of mice bearing subcutaneous or orthotopic PDAC tumors after treatment with vehicle (Ctrl) or DON. Data are expressed as mean ± SEM. KPC heterotopic (n=4), KPC orthotopic (n=8 and n=9), PaTu 8988T orthotopic (n=7), and 779E orthotopic (n=8 and n=7). b) Immunohistochemical staining of the apoptosis marker Cleaved Caspase 3 (CC3) in KPC and 779E orthotopic tumors treated with vehicle (Control) or DON (10mg/kg). Representative images are shown. Red scale bar 50 μm. c) Immunohistochemical staining of the proliferation marker pHis-H3 in 779E orthotopic tumors treated with vehicle (Control) or DON (10mg/kg). Representative images are shown. Scale bar 100 μm. Quantification of pHis-H3-positive nuclei/field is shown as mean ± SEM of n=4 tumors per group. Statistical significance was calculated using unpaired t-test in A and C. \**P*<0.05, \*\**P*<0.01

**Extended Data Fig. 2.**
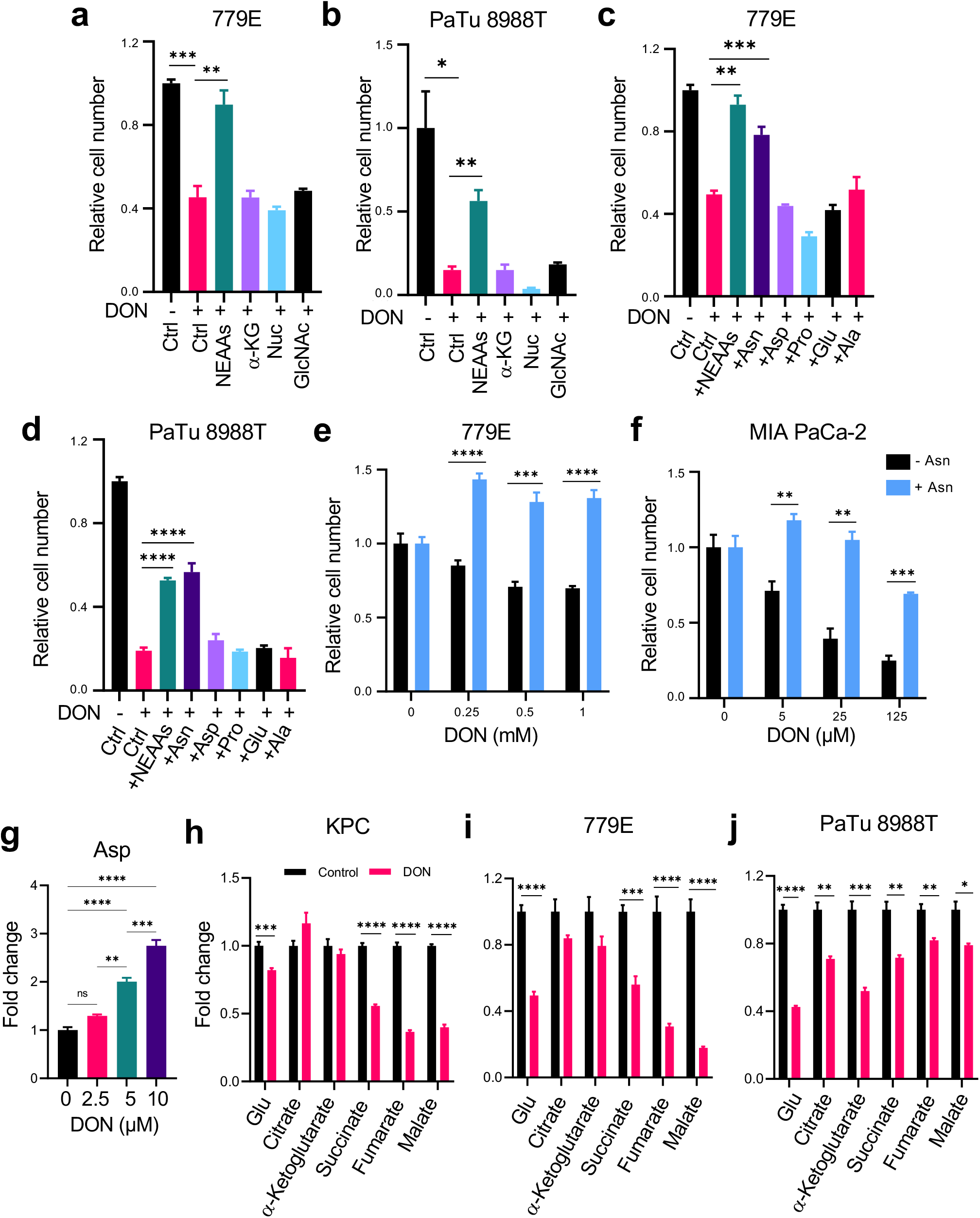
DON treatment affects polar metabolite pools but effects on cell fitness are only rescued by asparagine. a), b) Relative number of 779E (a) or PaTu 8988T (b) cells treated with or without DON (1mM or 10μM, respectively) with supplementation of the indicated metabolites for 24h. Cells were stained with crystal violet and stained area was quantified. Representative images are shown. Data are presented relative to untreated control for each condition and are representative of 3 independent experiments. Data are expressed as mean ± SEM of triplicate wells. c), d) Relative number of 779E (c) or PaTu 8988T (d) cells treated with or without DON (1mM or 10μM, respectively) supplemented with a cocktail of NEAAs or the indicated individual amino acids (0.1mM) for 24h. Representative images are shown. Data are presented relative to untreated control for each condition and are representative of 3 independent experiments. Data are expressed as mean ± SEM of triplicate wells. e), f) Relative number of 779E cells (e) or MIA PaCa-2 cells (f) treated with DON at the indicated concentrations with or without 0.1mM Asn supplementation. Data are presented relative to untreated control for each condition and are representative of 3 independent experiments. Data are expressed as mean ± SEM of triplicate wells. g) Quantification of intracellular aspartate (Asp) levels in PaTu 8988T cells treated with or without DON at the indicated concentrations for 24h. Data are presented relative to untreated control. Data are expressed as mean ± SEM of n=3 samples. h-j) Quantification of intracellular levels of the indicated TCA cycle metabolites in KPC (h), 779E (i) or PaTu 8988T (j) cells treated with vehicle control or DON (10μM, 10μM, and 1mM, respectively) for 24h. Data are presented relative to untreated control. Data are expressed as mean ± SEM of n=5 samples (h,i) and n=3 samples (j). Statistical significance was calculated using One-way ANOVA followed by Dunnet’s multiple comparisons test (a-d,g) or unpaired two-tailed Student’s t test in (e,f,h-j). \**P*<0.05, \*\**P*<0.01, \*\*\**P*<0.001, \*\*\*\**P*<0.0001.

**Extended Data Fig. 3.**
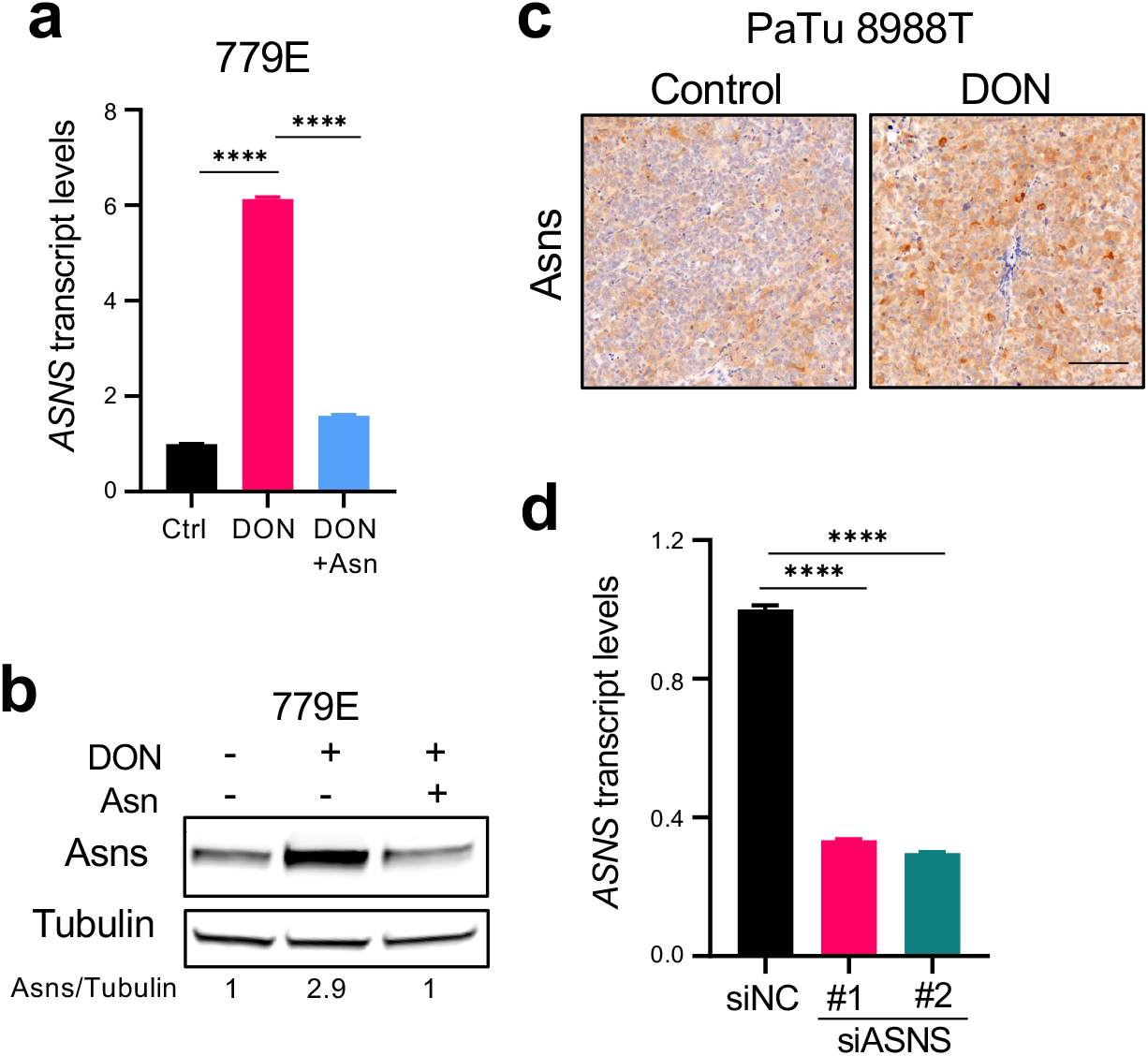
Determinations of ASNS expression levels using qPCR, immunohistochemistry, and immunoblotting. a) Relative *ASNS* mRNA expression levels as assessed by qPCR in 779E cells treated with 0.5mM DON with or without 0.1mM Asn supplementation for 24h. Data are presented relative to untreated control and are representative of 3 independent experiments. Data are expressed as mean ± SEM of n=3 replicates. b) Immunoblot assessing Asns protein expression in 779E cells treated with 0.5mM DON with or without 0.1mM Asn supplementation for 24h. Tubulin was used as a loading control. c) Immunohistochemical staining of Asns protein in PaTu 8988T orthotopic tumors treated with vehicle control (Control) or DON (5mg/kg). Representative images of n=4 mice per group are shown. Scale bar, 100μm. d) Relative *ASNS* mRNA expression levels in KPC cells as assessed by qPCR after transfection with non-targeting negative control siRNA (siNC) or two different hairpins targeting ASNS (siASNS#1 and siASNS#2) for 24h. Data are presented relative to siNC control and are representative of 3 independent experiments. Data are expressed as mean ± SEM of n=3 replicates. Statistical significance was calculated using One-way ANOVA followed by Tukey’s multiple comparisons test. \*\*\*\**P*<0.0001.

**Extended Data Fig. 4.**
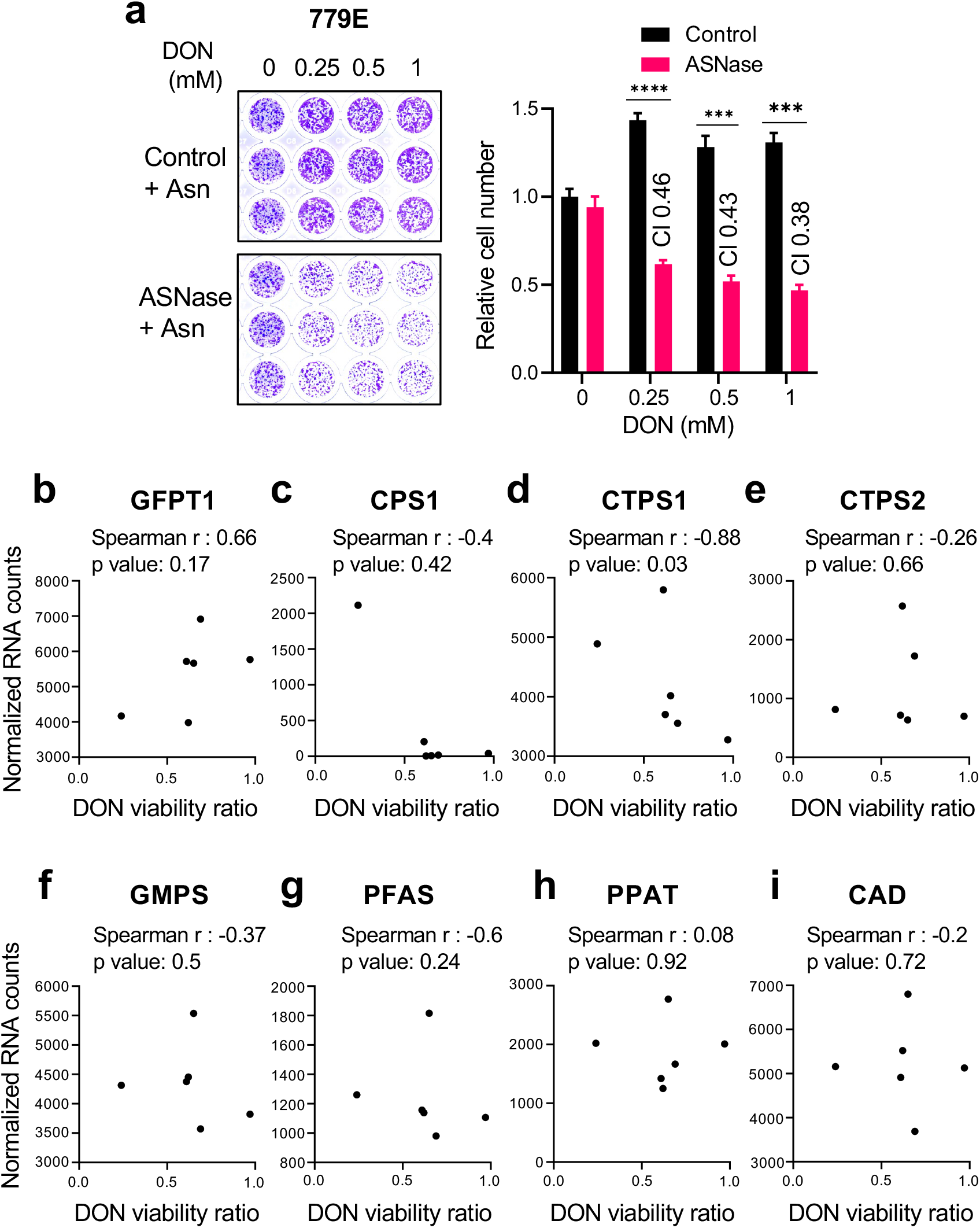
DON and ASNase co-treatment in 779E cells and correlation analyses for DON target genes in PDOs. a) Relative number of 779E cells treated with the indicated doses of DON in combination with ASNase (0.5U/ml) for 24h. Cells were stained with crystal violet and representative images are shown. Quantification of crystal violet staining is shown relative to untreated control and is representative of 3 independent experiments. The coefficient of drug interaction (CI) was calculated for each DON concentration (shown in graph). Data are expressed ± SEM of n=3 replicate wells. b-i) Correlation analysis between normalized gene expression of the indicated enzymes targeted by DON and DON viability ratio for the six PDOs assessed. Statistical significance was calculated using unpaired two-tailed Student’s t test (a) or Spearman correlation coefficient (b-j). \*\*\**P*<0.001, \*\*\*\**P*<0.0001.

**Extended Data Fig. 5.**
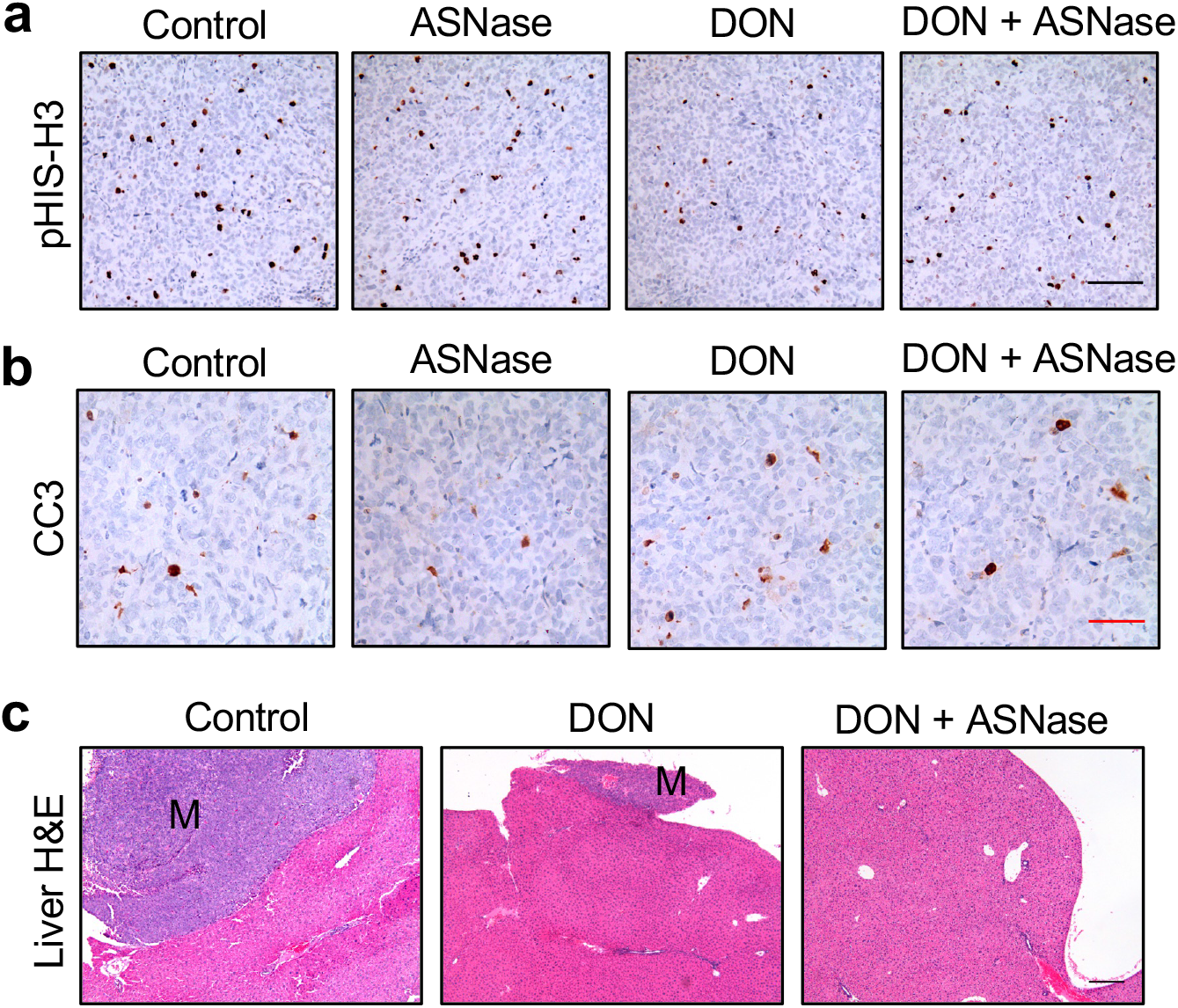
DON and ASNase co-treatment does not further impact cell death or proliferation in tumors, but does suppress liver metastases. a), b) Immunohistochemical staining of phospho-Histone H3 (pHis-H3; a) or cleaved Caspase 3 (CC3; b) in PaTu 8988T orthotopic tumors treated with vehicle (Control), ASNase (60U/mice), DON (5-10mg/kg), or a combination of DON + ASNase for 3 weeks. Representative images are shown. Scale bar 100 μm. c) Hematoxylin and Eosin (H&E) staining of liver sections from PaTu 8988T tumor-bearing mice treated with vehicle (Control), DON (5-10mg/kg), or a combination of DON + ASNase for 3 weeks. Metastases are indicated (M). Representative images are shown. Scale bar 200 μm.

